# Cortical dynamics underlying initiation of rapid steps with contrasting postural demands

**DOI:** 10.1101/2024.12.04.626758

**Authors:** Ilse Giesbers, Lucas Billen, Joris van der Cruijsen, Brian D. Corneil, Vivian Weerdesteyn

## Abstract

Our ability to flexibly initiate rapid visually-guided stepping movements can be measured in the form of express visuomotor responses (EVRs), which are short-latency (∼100ms), goal-directed bursts of lower-limb muscle activity. Interestingly, we previously demonstrated that recruitment of anticipatory postural adjustments (APAs) interacted with the subcortically-generated EVRs in the lower limb, suggesting context-dependent top-down modulation.

We investigated the associated cortical dynamics prior to and during rapid step initiation towards a salient visual target in twenty-one young, healthy individuals. We adopted two contrasting postural conditions by manipulating the stepping direction. Anterolateral steps involved low postural demands, whereas anteromedial stepping involved high postural demands. We recorded high-density EEG, surface electromyography from gluteus medius and ground-reaction forces.

Independent component analysis and time-frequency statistics revealed significant, yet relatively modest differences between conditions in preparatory cortical dynamics, most evidently in primary motor areas. Following target presentation, we observed stronger theta and alpha power enhancement in the supplementary motor area, and stronger alpha and beta power decrease in primary motor, parietal and occipital clusters during APA recruitment that preceded steps under high postural demands. We found no differences in (pre)frontal areas associated with the observed EVR suppression in the high postural demand condition.

Together, our findings point towards greater cortical involvement in step initiation under high postural demands as compared to more reflexive, stimulus-driven steps. This notion may be particularly relevant for populations where postural control is impaired by age or disease, as more cortical resources may need to be allocated during stepping.

## Introduction

Every day, we take thousands of steps without realizing the complex mechanisms underlying each step. Yet even before we start lifting the foot, goal-directed stepping movements are commonly preceded by so-called anticipatory postural adjustments (APAs). APAs involve coordinated activation of various muscles that induce a centre of mass shift towards the stance leg before unloading of the stepping leg. The APA is modulated according to the speed, size and direction of the upcoming movement to ensure balance during the step (Bancroft & Day, 2016; Inaba et al., 2020).

APAs can be initiated rapidly following stimulus presentation, but intriguingly, we recently demonstrated that a highly-salient visual target can evoke even faster visuomotor transformations that oppose the APA (Billen et al., 2023). Specifically, bursts of muscle activity in stance-leg gluteus medius occurred in a target-selective manner at average latencies of 107ms following left or right visual stimulus appearance, which facilitated rapid goal-directed stepping movements in the absence of an APA. These so-called Express visuomotor responses (EVRs; formerly called ‘stimulus-locked responses’) have been studied extensively in visually-guided upper-extremity movements (Corneil et al., 2004; Gu et al., 2016; Pruszynski et al., 2010) and are thought to be transmitted to the motor periphery along the subcortical tecto-reticulo-spinal pathway that originates in the superior colliculus (Boehnke & Munoz, 2008; Contemori et al., 2021b; Corneil et al., 2004; Corneil & Munoz, 2014; Pruszynski et al., 2010; Rezvani & Corneil, 2008).

While these visuomotor transformations are thought to be generated reflexively, the involved subcortical network is highly adaptive. For example, EVR output can be modulated by varying the temporal predictability of target appearance (Contemori et al., 2021a) or by changing the task instructions given to the participant (Contemori et al., 2023; Gu et al., 2016). Interestingly, stepping-related EVR expression was found to be modulated by the postural demands of the task (Billen et al., 2023). In this previous study, we manipulated the postural demands of the upcoming step by varying target location and stance width. In a low postural demand condition, participants stepped forward and outward towards the target from a narrow stance width. In this condition, EVRs were robustly expressed and facilitated rapid step initiation, while APAs were generally absent. In the high postural demand condition, participants stepped forward and inward from a wide stance width. Here, large APAs were present prior to foot-off, while EVRs were largely absent. Yet, whenever occasionally present in this condition, EVRs were followed by stronger, but delayed APAs with consequently longer stepping reaction times. These collective findings highlight the intricate interplay between EVR and APA expression, however, the neural mechanisms underlying their contextual modulation remain poorly understood.

Here, we aimed to gain insight into top-down modulation of fast goal-directed leg movements and postural control by studying cortical activity during step initiation towards a salient visual target. We used the same paradigm and postural demand manipulations as described previously (Billen et al., 2023) and additionally measured high-density electroencephalography (EEG) to uncover the dynamics of multiple cortical areas involved in proactive and reactive modulation of APA and EVR expression. We expected to find proactive differences (i.e. before stepping target appearance), because participants knew in advance (at least implicitly) if the postural demands of the upcoming step would be high or low. Reactive (i.e. after stepping target appearance) differences in cortical dynamics were expected between the postural demand conditions, reflecting contextual inhibition/facilitation of EVR and APA recruitment and ensuing differences in stepping behaviour.

## Materials and Methods

### Participants

A total of 21 healthy young individuals (14 female, age: 24.5 ± 2.0 years, range: 20-27) participated in this single-session study. Prospective participants were included in the study if they were aged between 18 and 35 years old and had a Body Mass Index of <25, to ensure high-quality EMG recordings. Exclusion criteria were visual, neurological or musculoskeletal disorders that may interfere with experimental task performance; behavioural problems interfering with compliance with the study protocol or pregnancy.

The study protocol was reviewed by the medical ethical committee (CMO Arnhem-Nijmegen, 2023-16109) and the study was conducted in accordance with the Declaration of Helsinki. All participants provided written informed consent before participation and were free to withdraw from the study at any time.

## Protocol

### Experimental set-up

The experiment was performed using a Gait Real-time Analysis Interactive Lab (GRAIL, Motek Medical, The Netherlands), which is situated at the Department of Rehabilitation at the Radboud University Medical Center. The set-up included an M-gait dual-belt treadmill with two embedded force plates (GRAIL, Motek Medical, The Netherlands) to measure ground reaction forces (sampled at 2000 Hz), a projector (Optoma, United Kingdom) to project the visual stimuli (see Figure 1B for the experimental set-up) and a photodiode (TSL250R-LF, TAOS, United States of America) to measure the exact moment of stepping target appearance.

**Figure 1.**
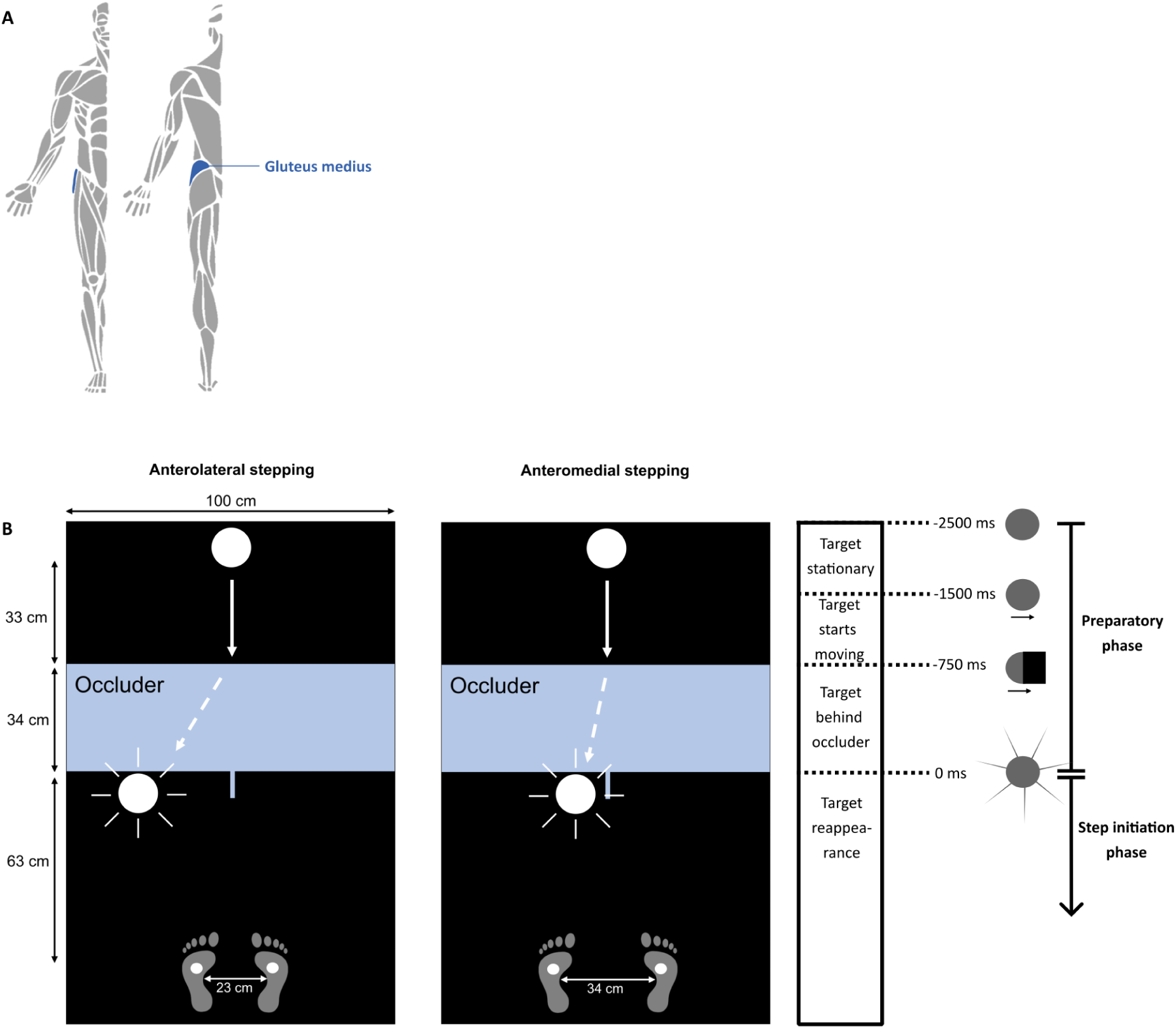
A: Location of the Gluteus Medius muscle. **B**: Schematic figure of the experimental paradigm (left) and time onset of events (right). Left: anterolateral stepping condition (foot width = 23 cm, distance target from midline = 29 cm). Right: anteromedial stepping condition (foot width = 34 cm, distance target from midline = 9 cm) along with time onset of the different phases in the experimental paradigm.

Further, electromyography (EMG) was recorded at 2000 Hz using a Wave Wireless electromyography system (Wave Wireless EMG, Cometa, Italy) from bilateral Gluteus Medius (GM) (Figure 1A) using Ag/AgCl surface electrodes. These were placed in accordance with the SENIAM guidelines (Hermens et al., 1999). A biosignal amplifier (REFA System, TMSi, The Netherlands) recorded high-density electroencephalography (EEG) from 126 scalp locations (Waveguard, ANT Neuro, The Netherlands) according to the five percent system (Oostenveld & Praamstra, 2001) at 2048 Hz without any filters, except for a built-in anti-aliasing filter at 552 Hz. Electrode impedances were kept below 10 kΩ. Using adhesive Ag/AgCl electrodes, the ground electrode was placed on the left mastoid and two electrodes were placed slightly above the nasion and at the outer cantus of the left eye to measure electrooculographic signals. Trials were started manually by the experimenter via the D-flow software (Motek Medical, the Netherlands). All reported measures (i.e. force plate data, EMG and EEG) were aligned to the moment of target appearance as detected by the photodiode.

### Experimental paradigm

Participants were instructed to divide their weight equally between their legs at the initial position indicated by the projection of small circles onto the treadmill (Figure 1B). At the start of a trial, a white, circular target was projected approximately 130 cm in front of the participant. After 1000 ms, the target started to move towards a projected occluder (750 ms) and disappeared behind it while the target retained its speed (750 ms) (i.e. *preparatory phase* (Figure 1B). Once the invisible target reached the base of the occluder, the target reappeared as a flash (for a duration of 48 ms) in front of the participant’s left or right leg and the participant stepped as fast as possible onto the target with the respective leg and placed the other leg alongside (i.e. *step initiation phase* (Figure 1B).

We adopted two different stepping conditions with contrasting balance demands. In the anterolateral stepping condition, participants stood in a narrow stance width (feet 23 cm apart) and stepped forward and outward towards an anterolateral target presented 29 cm from the middle line of the treadmill. In the anteromedial stepping condition, participants started at a wide stance width (feet 34 cm apart) and stepped forward and inward towards an anteromedial target, presented 9 cm from the middle line (Figure 1B).

The participants started with a few practice trials to familiarize themselves with the task. The experiment was divided into 4 blocks of 75 stepping trials. Two blocks consisted of exclusively anterolateral targets and the other two blocks consisted of exclusively anteromedial targets. The order of the blocks was counterbalanced between individuals and target side (left/right) was randomized in each trial.

## Data processing

All data (EMG, force plate and EEG) were analysed in MATLAB (Version 2020B, The Mathworks, Inc., USA) using custom scripts and were grouped based on target location and target side, which led to four different conditions (i.e. left-anterolateral, left-anteromedial, right-anterolateral, right-anteromedial). Incorrect trials were excluded from all analyses (EEG, EMG and force plate) and were defined as trials in which participants stepped towards the wrong direction or initiated a stepping movement with the contralateral foot. Further, trials were excluded from the analysis if participants showed a stepping reaction time of less than 200 milliseconds, as such response times indicate a person would have guessed the stepping side and already initiated the movement before trial onset. Trials were also excluded from analysis if the participants exhibited reaction times slower than 1000 milliseconds.

### EMG data processing

The raw EMG signals were band-pass filtered between 20 Hz and 450 Hz and subsequently rectified and low-passed filtered at 150 Hz using a second-order Butterworth filter. Like prior studies studying EVRs (Billen et al., 2023; Gu et al., 2016; Kozak et al., 2020), a receiver operating characteristic (ROC) analysis was performed to determine the presence and latency of the EVRs. EVR latency was defined at the time at which the AUC surpassed the discrimination threshold (0.6) and remained above this threshold for 16 out of 20 consecutive samples within the pre-defined EVR epoch of 100 to 140 milliseconds after target reappearance.

Regardless of whether an EVR was found, the response magnitude in the EVR window was computed for every condition within every participant. The mean EMG activity of the 20 ms window, which was centered around the maximum EMG activity during the EVR epoch (100-140 ms), was determined on a single trial basis. Magnitudes were then normalized to the median peak EMG activity during anteromedial stepping (in the interval from 140 ms to foot-off). The average EMG magnitudes for each condition were then calculated.

### Force plate data processing

The raw force plate data was used to determine stepping reaction times (i.e. the time between target reappearance and the moment of foot-off of the stepping leg). Average stepping reaction times were calculated for each condition and stepping side. APA onset was calculated based on the stepping-side and stance-side force plate data using a time-series ROC analysis, whereby the onset was defined as the time at which the AUC surpassed the discrimination threshold (0.6) for 8 out of 10 consecutive samples within the 100-300ms window following target reappearance. APA magnitude was defined as the baseline-corrected difference between the mean maximum vertical ground reaction force component (Fz) underneath the stepping leg and the corresponding vertical ground reaction force underneath the stance leg in the interval from 140ms after target appearance (i.e., the end of the EVR window) and foot off, normalized to percent total body weight (%BW). APA magnitude was calculated regardless of whether the ROC analysis identified a significant APA.

### EEG data preprocessing

For analysing the EEG data, we used custom scripts and incorporating functions from EEGLAB (Version 2023.0; Delorme & Makeig, 2004a). EEG data were band-pass filtered between 1 Hz and 200 Hz using a 4^th^-order Butterworth filter FIR with a zero-phase shift. Additional notch filters were applied to reduce the line noise of 50 Hz and its harmonics of 100 and 150 Hz. For each EEG channel, z-scores were calculated from the sum of each channel signal over time. Channels with z-scores >1.96 were considered outliers and were rejected. Furthermore, channels with a probability and kurtosis measure exceeding 6 standard deviations from the mean were rejected. Data were visually inspected for flat channels and were also removed. On average, a total of 20 channels (*SD* = 11.2) were removed per participant and the remaining EEG channels were re-referenced to the common average. Next, the EEG data were segmented in epochs of -3.5 to +1 seconds relative to the target reappearance. Epochs consisting of wrong, too slow or too fast steps (for more information, refer to section 2.3) were removed. Next, automated artefact rejection was performed by calculating the joint probability of all channels at each timepoint (local and global threshold of 6) and rejecting epochs with exceeding a probability of 6 times the standard deviation. On average, 10% of all trials were removed (*M* = 31, *SD* = 21.7) for each participant.

## Source separation

Subsequently, data were downsampled to 256 Hz to speed up data processing. An Infomax Independent Component Analysis (ICA) was performed on the preprocessed EEG data to isolate and extract contributions of different underlying cortical regions giving rise to the EEG signal, while separating the influence of artefactual signals, such as eye movements and muscle activity. This approach is in line with other studies on cortical dynamics during whole-body movements (Gwin et al., 2011; Solis-Escalante et al., 2019; Wagner et al., 2016).

Next, locations and orientations of the underlying neural activity were estimated by fitting an equivalent current dipole to the scalp projection of each Independent Component (IC) using the DIPFIT toolbox (Delorme & Makeig, 2004; Oostendorp & Oosterom, 1989). A standardized three-shell boundary element head model consisting of standard scalp, skull and brain conductivities (0.33 S/m, 0.0041 S/m and 0.33 S/m, respectively; Oostenveld et al., 2003) and standard electrode positions were used to find the location and orientation of the cortical sources. Independent components related to eye movements and muscle artifacts were identified through visual inspection of their power spectra, scalp topography and location of their associated equivalent current dipole. Using EEGLAB’s *IClabel,* we identified ICs related to brain activity, showing typical EEG power spectra and equivalent current dipole inside the head. ICs with low residual variance (< 15%) were further analysed. This resulted in an average of 10 components per participant (SD = 3.73).

## Event-related spectral perturbations

Oscillatory dynamics during the preparation and initiation of a stepping movement were quantified using event-related spectral perturbations (ERSPs), which provide information about how the spectral power is modulated over time relative to an event. Frequency bands were defined as follows: theta (3-8 Hz), alpha (8-13 Hz) and beta (13-30 Hz) (Liu et al., 2024; Zhao et al., 2022). The beta band was subdivided in low beta (beta I: 13 – 18 Hz) and high beta (beta II: 18-30 Hz) (Solis-Escalante et al., 2019). We precomputed component measures using EEGLAB’s *pop_precomp* function, using Morlet wavelets (frequencies between 3 and 30 Hz, with 3 cycles at the lowest frequency and increasing cycles linearly by 0.8 with each step, baseline between –3500 and –2700).

Subsequently, ICs of each participant were clustered using principal component analysis with k-means algorithm and were clustered based on similarities in their spatial location of their associated current dipole. The number of clusters was set as the average number of ICs for each participant (n=10). ICs that were more than three standard deviations away from the cluster centroid were identified as outliers and removed. If clusters held more ICs from one participant, we selected the IC with the lowest residual variance to prevent artificially inflating the sample size. Clusters that contained ICs of more than half of the participants were further analysed (≥ 11). Further, we computed power spectral densities of each cluster to identify clusters showing spectral changes compared to baseline.

## Statistical analysis

### Behavioural measures

After processing the EMG data and performing an ROC analysis, average EVR onset times and EVR magnitudes were calculated for each condition and stepping side. Paired samples t-tests were performed to study differences between stepping conditions and sides.

Similarly, to the EMG data, APA latencies, magnitudes and stepping reaction times for each stepping condition and stepping side were calculated based on the force plate data. Paired samples t-tests were performed to study differences between stepping conditions and sides.

### Cortical dynamics

For each cluster that met the requirements, we evaluated differences in cortical activity between anterolateral and anteromedial stepping. We averaged the time-frequency maps over all ICs in a given cluster, for each stepping condition separately. Next, we subtracted the average ERSP of the anterolateral condition from the average ERSP of the anteromedial condition, to create contrast maps which allowed us to visualize and interpret differences in cortical activity between stepping conditions. We used permutation-based testing (Maris & Oostenveld, 2007) to evaluate differences across stepping conditions (2000 iterations, α = .05), as was previously done in other studies using EEG measures during whole-body movements (Solis-Escalante et al., 2019). We report both False Discovery Rate (FDR)-corrected (Benjamini & Hochberg, 1995) as well as uncorrected differences between stepping conditions, to avoid being overly conservative and thereby resulting in an increased risk of false negatives (type 2 error).

## Results

Any differences between stepping side (left/right) and muscles (left/right GM) in behavioural and EMG-related outcomes were not significant. We therefore averaged all outcomes across sides.

## Behavioural results

Overall, participants’ error rates were low, on average 2% (*M* = 4.9%, *SD* = 4.6%) in all 300 trials combined. Participants made more mistakes in the anteromedial stepping condition (*M* = 4.3%, *SD* = 3.5%) compared to the anterolateral condition (*M* = 1.1%, *SD* = 1.7%).

Furthermore, participants showed an average of 4 trials (*M* = 4.1, *SD* = 6.0) with reaction times that were too early (i.e. faster than 200 milliseconds) across all 300 trials. No trials with reaction times slower than 1000 milliseconds were detected.

## EMG and force plate data in a representative participant

To demonstrate the timeline of behavioural events, Figure 2 shows EMG and force plate data of left GM when it is on the stance side (dark blue) and when it is on the stepping side (light blue) during anterolateral stepping (top row) and anteromedial stepping (bottom row) for a representative participant. In anterolateral stepping, *stance side* GM (dark blue traces) rapidly becomes active within 120ms after target onset, whereas the *stepping side* GM remains relatively silent (first column). The second column shows trial-by-trial EMG traces in the *stance* leg sorted by stepping reaction time, marked by white dots. This plot illustrates two distinct bursts of muscle activity, with the first occurring approximately 110 ms after target reappearance. This initial burst of activity is the EVR, as it is time-locked to target reappearance and remains independent of the subsequent stepping reaction time. The second burst corresponds to voluntary muscle activation driving the stepping movement. The third column reveals left GM activation when it is on the *stepping* side, demonstrating its relative silence throughout the trial. The fourth column depicts the average force plate data for both the stepping and stance leg. At ∼150ms, the vertical forces immediately decrease on the stepping side with a concurrent increase underneath the stance leg, indicating the absence of the initial weight shift that typically signifies the execution of an Anticipatory Postural Adjustment (APA).

**Figure 2.**
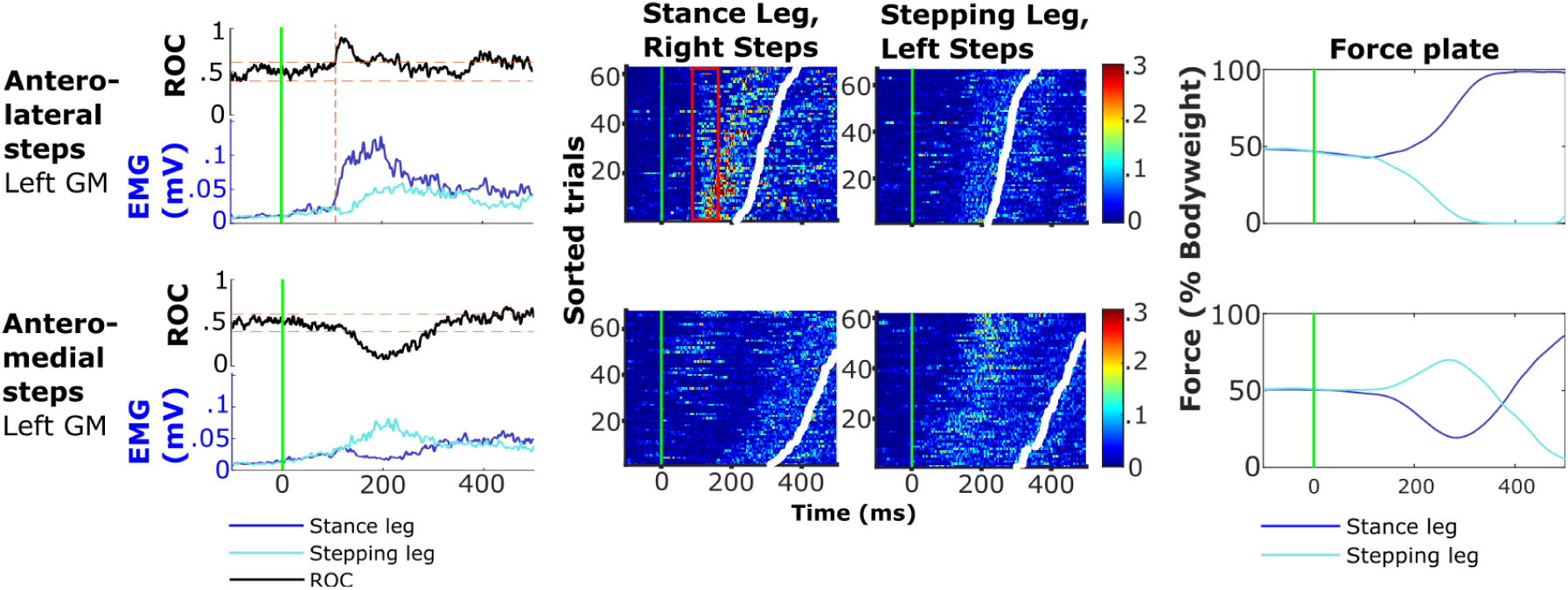
Left GM muscle activity, time-series ROC analysis and force plate data of an exemplar participant for anterolateral stepping condition (top row) and anteromedial stepping condition (bottom row). Column 1: ROC analysis (black line) and mean EMG activity in mV for when recorded from stance (dark blue) or stepping leg (light blue), and EVR latency (dashed red vertical line). Column 2 and 3: Trial-by-trial EMG in the left stance leg GM (column 2) and stepping leg GM (column 3). Column 4: Mean vertical force (relative to percentage of the participant’s body weight) exerted by stance leg (dark blue) and stepping leg (light blue); the initial increase in stepping leg force in the anteromedial condition is the APA.

In the anteromedial stepping condition (bottom row), EMG activity and force plate data are distinctly different. Here, the *stance leg* GM remains relatively silent until shortly before step onset, as indicated by the first column. Yet, inspection of the individual EMG traces of the *stance leg* GM (second column) reveals occasional EVR-type activity that is primarily present on the slower half of trials, which is then promptly suppressed. The individual EMG traces in the *stepping leg* (third column) shows pronounced muscle activity in the stepping leg GM between ∼150-250 ms. This burst of muscle activity is temporally linked to APA execution, as shown by the increase in vertical force underneath the stepping leg (fourth column), which pushes the CoM towards the stance side prior to lifting the foot off the ground at ∼500ms.

## Robust EVRs during anterolateral stepping

In line with previous findings (Billen et al., 2023, 2024), we observed robust EVRs in 18 of the 21 participants during anterolateral stepping, for both left GM (16/21) and right GM (17/21) with an average EVR latency of 117 ms (SD = 6 ms). None of the 21 participants exhibited consistent EVRs in the anteromedial stepping condition. In line with observations from the visual inspection of the representative participant and consistent with the absence of detectable EVRs in anteromedial stepping, response magnitudes within the EVR window were significantly higher in anterolateral stepping (*M* = .10, *SD* = .05) compared to anteromedial stepping (*M* = .05, *SD* = .01; *t(20)=* 5.6, *p* < .001, *Hedges g* = 1.42) (see Figure 3A).

**Figure 3.**
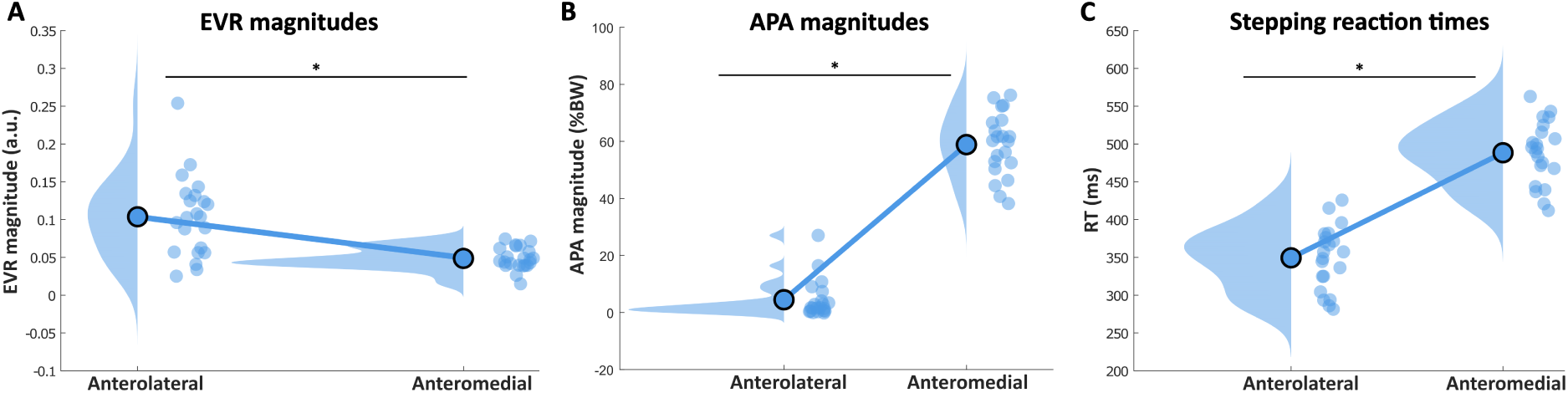
A-C: EMG and behavioural outcomes of all participants for anterolateral and anteromedial stepping. The dots indicate individual averages, averaged across left and right steps. The density plots indicate the distribution of the data. Asterisks indicate significant differences (p < .05). For A and B, EVR and APA magnitudes are shown for all participants, independent of whether the ROC analysis detected a significant EVR/APA.

In order to identify the occasional expression of EVRs in the anteromedial condition, we split the trials in the anteromedial stepping condition into a fast- and a slow-RT half and separately performed the time-series ROC analyses within the EVR window on the two subsets of trials. Replicating our previous findings (Billen et al., 2023), EVRs were present on the slow half of trials in 17 out of the 21 participants in at least one GM (4 with bilateral EVRs).

## Prominent APAs and slow reaction times in anteromedial stepping

All participants produced robust and strong APAs in the anteromedial stepping condition, whereas only a small number of participants produced APAs during anterolateral stepping (4/21 during leftward steps, 6/21 during rightward steps). In anteromedial stepping, APAs were initiated at 162ms (SD = 13 ms) after target reappearance, which did not differ significantly from APA onset times in anterolateral stepping (M = 170 ms, SD = 23 ms; *t*(9) = 1.4, *p* = 0.20). APA magnitudes were significantly larger in anteromedial stepping (*M* = 0.59 %BW, *SD* = 0.11 %BW) compared to anterolateral stepping (*M* = 0.04 %BW, *SD* = 0.07 %BW; *t*(20) = -24.5; *p* < .001; *Hedge’s g* = -5.9; see Figure 3B).

Overall, stepping reaction times were significantly faster for anterolateral steps (*M* = 349 ms, *SD* = 43 ms) than anteromedial steps (*M* = 488 ms, *SD* = 43 ms; *t*(20) = -23.1, *p* < .001, *Hedges g* = -3.35, see Figure 3C).

## Task-related cortical dynamics

Clustering the independent components using k-means algorithm led to 10 IC clusters. Two clusters were not further considered due to the absence of spectral changes compared to baseline (left temporal cluster) and containing ICs from less than half of the participants (right temporal cluster). 8 IC clusters were further considered. Table 1 shows the Talairach coordinates of the cluster centroids, which provide an estimation of the location of the actual cortical sources, constrained by the spatial resolution of the source localization methods (standard electrode positions and standard head model). We found one frontal cluster (located in the anterior cingulate cortex). We found three clusters in cortical motor areas: a midline central cluster, which was located in the supplementary motor area and two lateralised clusters, located in the left and right primary motor cortex. We also found three parietal clusters in the posterior cingulate (ventral posterior cingulate), the midline parietal (visuomotor area) and the posterior parietal cortex (angular gyrus). We further found an occipital cluster, which was located in the left visual association area. Figure 4 displays the spatial location of each independent components for each cluster (first row) including the cluster centroid, as well as the spatial location of the cluster centroids only (second row).

**Figure 4.**
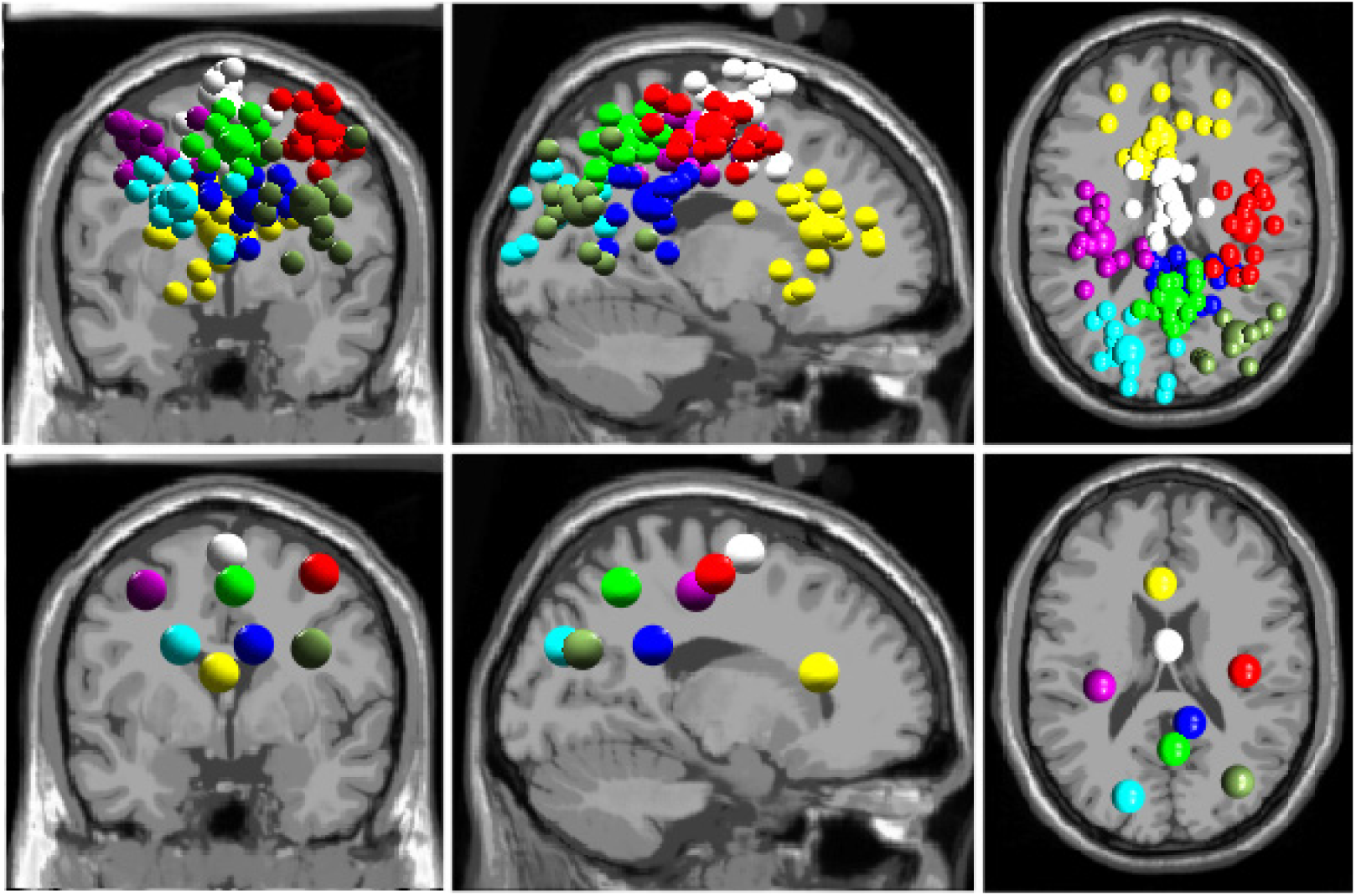
Dipole locations for all participants (top row) and cluster centroids (bottom row) in sagittal (left), axial (middle) and coronal (right) plane. Cluster centroids are displayed in a larger size and colours correspond to their respective clusters. We identified 8 relevant clusters, anterior cingulate (yellow), primary motor area left (purple) and right (red), midline central (white), posterior cingulate (blue), midline parietal (lime), posterior parietal (olive) and left occipital (aqua).

**Table 1.** Estimated location of clusters centroids.

| Lobe | IC Cluster | ICs | Talarach coordinates (x, y, z) | Cortical location | Brodmann area | Colour |
| --- | --- | --- | --- | --- | --- | --- |
| Frontal cortex | Anterior cingulate | 18 | -3, 25, 14 | Anterior cingulate (L) | BA 24, range = 1* | Yellow |
| Motor cortex | Primary motor area (L) | 19 | -33, -22, 47 | Primary motor area (L) | BA 4 | Purple |
|  | Primary motor area (R) | 19 | 37, -14, 53 | Primary motor area (R) | BA 4, range = 1* | Red |
|  | Midline central | 17 | 0, -2, 60 | Supplementary motor area | BA 6 | White |
| Parietal cortex | Posterior cingulate | 14 | 11, -40, 27 | Ventral posterior cingulate (R) | BA 23 | Blue |
|  | Midline parietal | 15 | 3, -51, 49 | Visuomotor area (R) | BA 7 | Lime |
|  | Posterior parietal | 11 | 34, -68, 27 | Angular gyrus (R) | BA 39 | Olive |
| Occipital cortex | Left occipital | 12 | -19, -75, 29 | Left visual association area | BA 19 | Aqua |
| Temporal cortex | Left temporal<sup>1</sup> | 12 | -46, -16, 7 | Left primary auditory | BA 41 | - |
|  | Right temporal<sup>2</sup> | 9 | 39, 1, 10 | Right insula | BA 13 | - |

\*Outside of defined BAs, nearest grey matter is noted. Range = 1 means within 5x5x5 mm search range.
<sup>1</sup> Did not display significant spectral changes compared to baseline.
<sup>2</sup> Did not meet requirements of number ICs $\geq 11$ .

## Frontal cortex

In the anterior cingulate cluster, we did not find noteworthy spectral changes compared to baseline during step preparation (Figure 5). Following target reappearance, we observed an increase in the theta- and alpha bands, that did not significantly differ between conditions.

**Figure 5.**
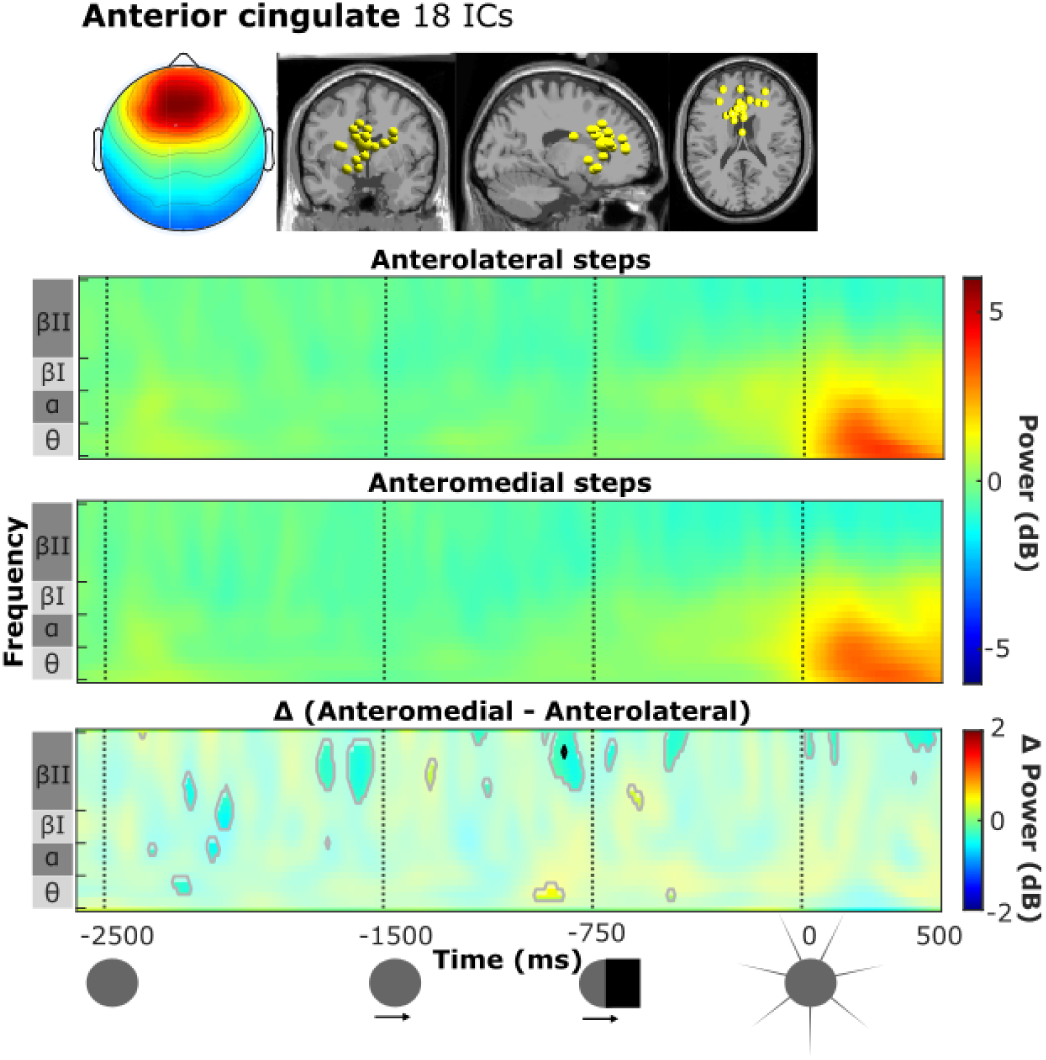
Scalp map and locations of equivalent current dipoles (grey underlay), ERSPs for anterolateral (first row) and anteromedial steps (second row) and contrast plot (third row) for the anterior cingulate cluster. ERSPs averaged time-frequency maps across participants for the anterolateral (top row) and anteromedial (second row) stepping conditions, and the contrast plot (third row) indicating differences between conditions (anteromedial minus anterolateral). Frequency bands are indicated on the y-axis. The vertical dashed lines in the time-frequency plot correspond to trial-related events (icons below) and indicate first target presentation (t = -2500 ms), onset of target movement (t = -1500 ms), disappearance of target behind occluder (t = -750 ms) and target reappearance (t = 0 ms). Time-frequency maps show a decrease (blue) and increase (red) in mean power, relative to the baseline. In the contrast plot, the non-significant differences (α = .05) are overlayed by a white transparent mask. Non-FDR corrected significant differences are indicated by a grey contour, FDR corrected significant differences are indicated by a black contour.

## Supplementary and primary motor cortical areas

During the preparation of an upcoming step, time-frequency analysis of the midline central cluster (Figure 6) indicated a beta II power decrease that started shortly after the onset of target motion and lasted until target reappearance, yet with little differences between conditions. Around target reappearance, a long-lasting alpha/theta power increase occurred, which was significantly stronger during the initiation of an anteromedial compared to an anterolateral step. Step initiation was also accompanied by a beta II power decrease. A brief, yet noteworthy significant difference occurred in the beta I frequency band, where participants displayed stronger power suppression during anteromedial stepping compared to anterolateral stepping.

**Figure 6.**
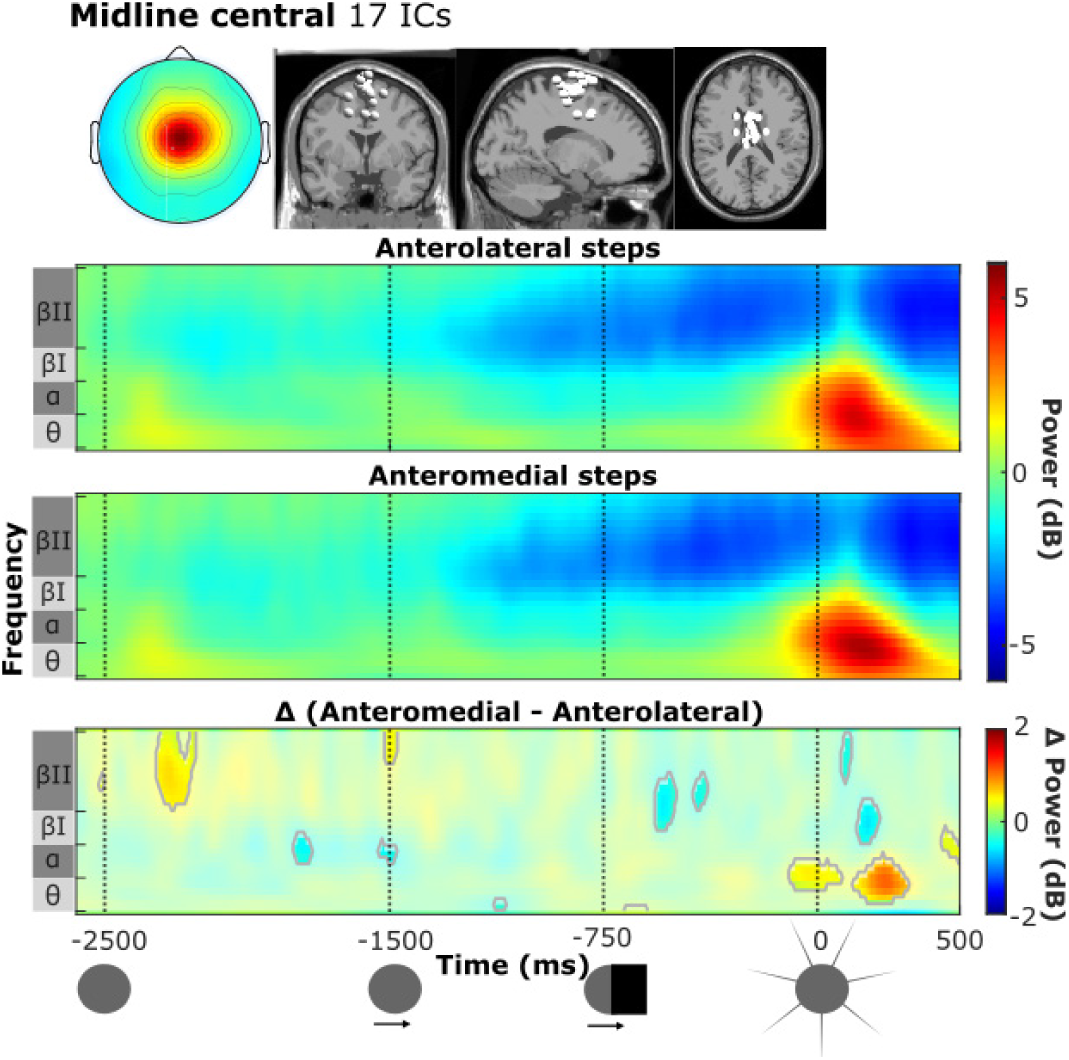
Scalp map and locations of equivalent current dipoles (grey underlay), ERSPs for anterolateral (first row) and anteromedial steps (second row) and contrast plot (third row) for the midline central cluster. Same format as Figure 5.

Time-frequency analysis of the left and right primary motor cortex presented a similar oscillatory pattern compared to baseline and similar pattern of significant differences between stepping conditions and we therefore only report the results of the left primary motor cortex (please refer to Appendix Figure 1A for ERSPs of the right primary motor cortex). The M1 (Figure 7A) presented a strong low-alpha (8-10 Hz) power suppression throughout the preparatory phase. Significant differences between stepping conditions mainly occurred in the high-alpha band during the stationary and early moving target phase, where participants displayed a modest power increase relative to baseline during preparation of an anterolateral step and a modest power decrease compared to baseline during the preparation of an anteromedial step. Around target reappearance, high alpha and beta I rhythms also showed significantly more suppression in preparation of an anteromedial step.

**Figure 7.**
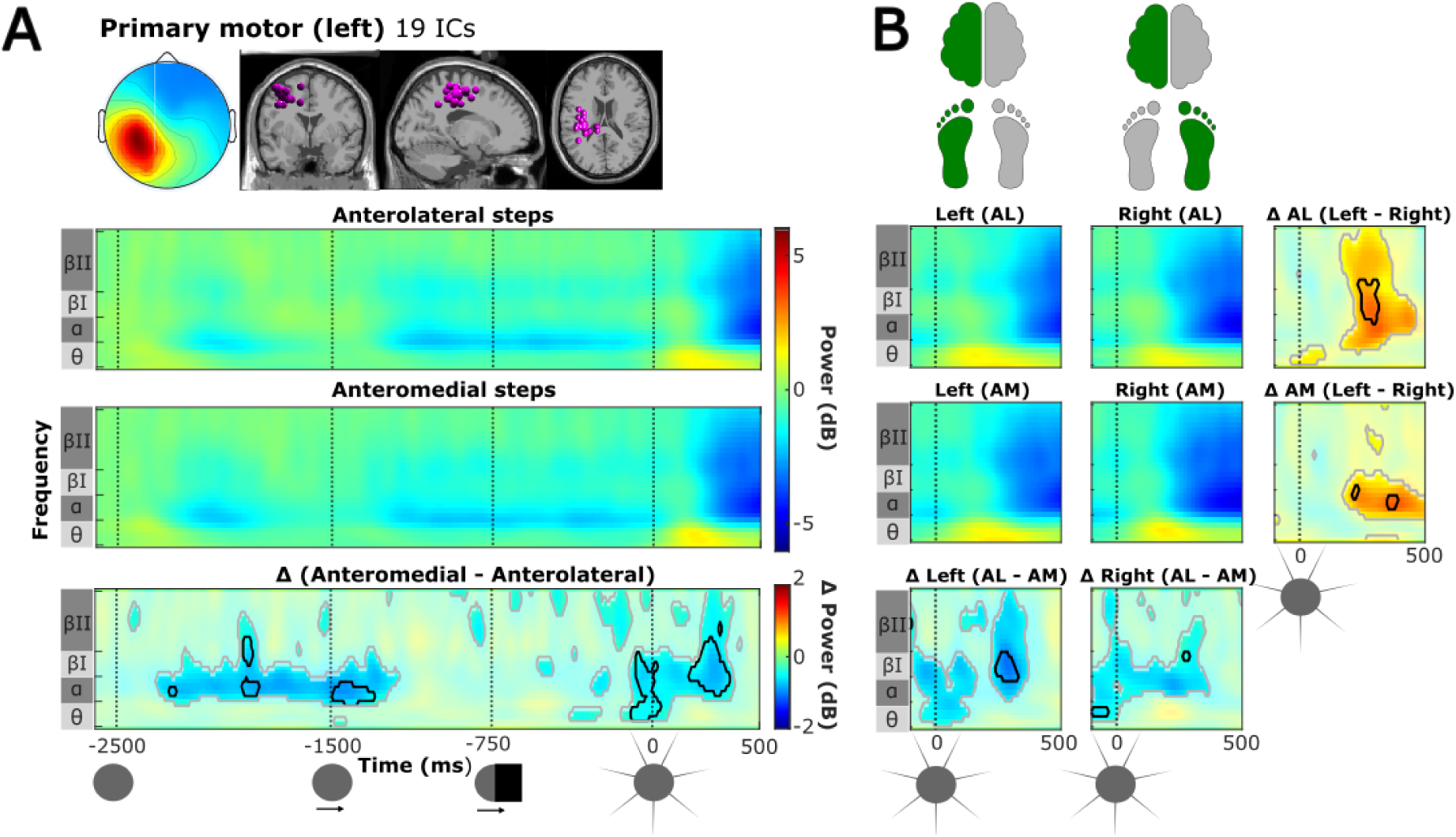
**A**. Scalp map and locations of equivalent current dipoles (grey underlay) for the left motor cluster, ERSPs for anterolateral (first row) and anteromedial steps (second row) and contrast plot (third row) for the left primary motor cluster. Same format as Figure 5. **B.** ERSPs during the step initiation phase (from target reappearance t = 0) for left (first column) and right steps (second column) during anterolateral stepping (first row) and anteromedial stepping (second row) for the left primary motor cluster. The contrast maps in the third column display the lateralized differences during anterolateral stepping (first row) and anteromedial stepping (second row). The contrast maps in the third row display differences in stepping conditions during left steps (first column) and right steps (second column).

During step initiation, the M1 displayed a weak power increase in theta and a strong broadband power decrease over alpha, beta I and beta II bands. This power decrease initially differed significantly between conditions, where participants showed stronger broadband power decrease in alpha, beta I and beta II-bands during anteromedial stepping compared to anterolateral stepping.

## Lateralized differences during anterolateral and anteromedial stepping in motor cortex

We observed strong lateralized activity in both the right and left M1 during the step initiation phase. Since the results of right and left M1 were essentially mirrored, we only report results from left M1 for conciseness (Figure 7B; see Appendix Figure 1B for right M1 results). During anterolateral step initiation, the left M1 displayed significantly greater power reductions in the alpha and beta bands for steps with the (contralateral) right leg compared to the left leg (Figure 7B, top row). These lateralized differences were less pronounced and largely restricted to the alpha band during anteromedial step initiation (Figure 7B, second row).

## Parietal and occipital cortex

Overall, time frequency analysis of the parietal and occipital clusters displayed a similar oscillatory pattern during preparation of the upcoming step. Therefore, we here show the ERSPs and contrast map of the cluster with greatest contrast between conditions (the posterior cingulate cluster; Figure 8). Please refer to the Appendix for the ERSPs and contrast maps of the midline parietal, posterior parietal and occipital cluster (Figures 1-4).

**Figure 8.**
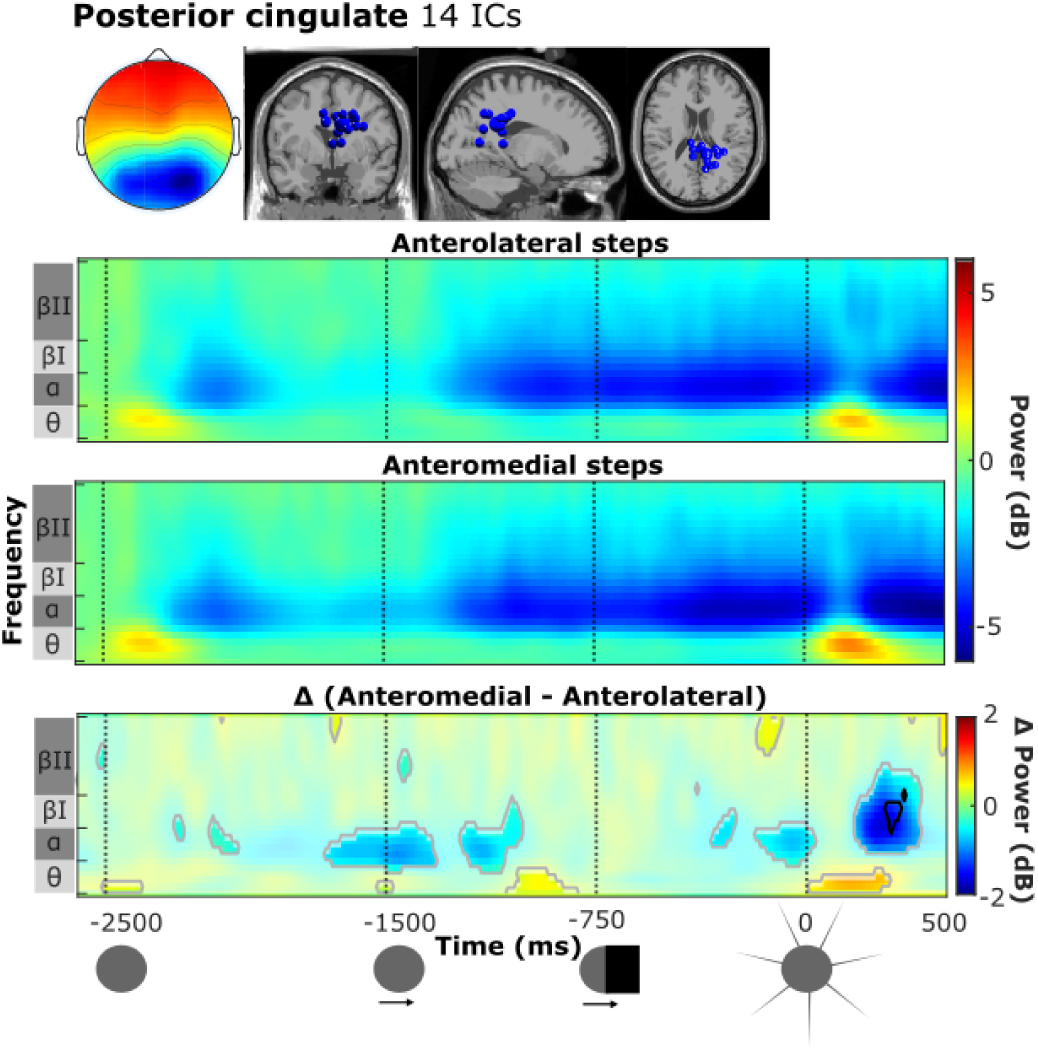
Scalp map and locations of equivalent current dipoles (grey underlay), ERSPs for anterolateral (first row) and anteromedial steps (second row) and contrast plot (third row) for the posterior cingulate cluster. For detailed description, see Figure 5.

The preparatory oscillatory pattern consisted of an alpha/beta I power decrease following initial target presentation, followed by a strong, continuous alpha/beta I during the real and implied motion of the target. Sparse significant differences between stepping conditions were observed in the alpha band in the posterior cingulate (Figure 8) and occipital cluster (Appendix Figure 4), where stronger alpha power suppression was found during the preparation of an anteromedial step compared to an anterolateral step.

During step initiation, we observed a transient theta power enhancement, followed by a strong alpha/beta I power decrease over all parietal and the occipital clusters. The theta enhancement was significantly stronger during anteromedial stepping in the posterior cingulate cluster (Figure 8), and to a lesser degree in the midline parietal cluster (Appendix, Figure 3). Additionally, strong alpha/beta I power decreases were observed in all four parietal and occipital clusters. This power decrease was strongest in the posterior cingulate and occipital cluster, and significantly more so in anteromedial compared to anterolateral stepping.

## Discussion

The current study investigated differences in oscillatory activity from multiple cortical areas during the preparation and initiation of rapid stepping movements between a low-postural demand (anterolateral stepping) and a high-postural demand condition (anteromedial stepping). Replicating our previous findings, we observed a reciprocal relationship between EVRs and APAs: when postural demands were low, EVRs were present in most participants and APAs were mostly absent, but when postural demands were high, EVRs were largely suppressed and instead APAs were strongly expressed (Billen et al., 2023). Cortical dynamics were evaluated for two separate phases within the task, namely the preparatory phase and the step initiation phase. During the preparatory phase (i.e. from initial target presentation until target reappearance), we found significant differences in cortical dynamics between anterolateral and anteromedial stepping conditions that were most evident in primary motor areas. During the step initiation phase (i.e. from target reappearance until the foot was lifted to start the stepping movement), a transient theta and alpha power increase was observed in multiple clusters, most strongly in the SMA. Here, this alpha/theta power increase was significantly stronger in the anteromedial compared to the anterolateral stepping condition. Shortly following the theta/alpha power increase, there was a widespread alpha and beta power decrease, which was significantly stronger during anteromedial step initiation in parietal, occipital and primary motor clusters.

## Cortical dynamics in preparation of step initiation with distinct postural demands

Before discussing the observed differences in preparatory cortical dynamics, we first reiterate what (motor) actions can already be planned ahead, regardless of stepping direction. Prior to target reappearance, the participant knows 1) the temporal dynamics of the task, i.e. *when* the step will have to be initiated and 2), due to the blocked nature of this study, what the postural demands of the upcoming step are, i.e. whether an APA needs to be expressed or not. Importantly, it is unknown *what* the exact stepping movement will be, as stepping side and, consequently, the direction of the APA and the stepping leg are only determined upon target reappearance.

In line with the known involvement of the supplementary and primary motor areas in APA control (Bolzoni et al., 2015; Brugger et al., 2020, 2021; Gwin et al., 2011; Jacobs et al., 2009; Ng et al., 2011, 2013), we observed modulations in the alpha and beta rhythms relative to baseline during the second half of the preparatory phase (i.e. until target reappearance). In the supplementary motor area, we found a prominent (high) beta decrease, but this was independent of the postural condition and thus argues against its proactive involvement in upregulation of postural circuitries involved in APA expression. The general consensus about (motor cortical) alpha and beta oscillations is that they are inhibitory in nature, where a power decrease reflects a gradual release from inhibition, or in other words, stronger cortical activation (Kilavik et al., 2013; Pfurtscheller, 2006; Pfurtscheller & Lopes da Silva, 1999). Beta power suppression is commonly observed both in anticipation of and during execution of a movement, followed by a power increase after movement execution (Engel & Fries, 2010; Kilavik et al., 2013). A previous EEG study reported a similar centrally-located beta power decrease starting around 2 seconds before voluntary step onset, coinciding with the preparatory phase of step initiation (Varghese et al., 2016). Unlike the study of Varghese and colleagues (2016), our paradigm did not allow directional APA planning (i.e. which side to step towards, as this was only evident for the participant upon target reappearance). Thus, SMA beta dynamics prior to step initiation may reflect the planning and temporal coordination of global motor actions that need to be performed upon target reappearance, irrespective of the postural demands and target side. We speculate that this global action may include the forward thrust of the CoM required for step initiation, as early (i.e. within 120 ms following target reappearance) bilateral activation of tibialis anterior in anteromedial as well as anterolateral steps was previously shown using the same task (Billen et al., 2023). Such non-lateralized muscle activation can therefore be planned without knowing the direction of the upcoming stepping movement.

In the primary motor cortex (both left and right cluster), we did observe significant differences in preparatory dynamics in the high-alpha and low beta band between postural conditions. During the preparation of an anterolateral step, participants displayed a modest power *increase* relative to baseline, whereas participants displayed a modest power *decrease* during the preparation of an anteromedial step. This observation may indicate a preparatory shift to overall greater motor cortical involvement in steps that require APA recruitment, as compared to the more reflexive generation of APA-less anterolateral steps. Neurons in M1 have direct corticospinal projections, but also project onto reticulospinal pathways (Fisher et al., 2021). As such, the observed alpha and beta suppression in primary motor cortex may be involved in upregulation of spinal as well as supraspinal postural circuits in preparation of APA generation.

Interestingly, parietal areas also exhibited differences in preparatory cortical dynamics, with participants showing a significantly stronger suppression mainly in the alpha-band during the preparation of an anteromedial step compared to an anterolateral step. From a cognitive perspective, the posterior parietal cortex is thought to act as a visuomotor controller, making computations based on proprioceptive, visual and attentional resources and may set the selection, preparation and execution of motor actions in motion by reciprocally interacting with the premotor areas (Goodale, 2011; Rilk et al., 2011; Wise et al., 1997). Thus, we speculate that the decreased preparatory alpha band power may point at increased visual-spatial attention toward the target under high postural demand, perhaps recruiting necessary premotor or visuomotor areas where attentional signals may be the modulators.

In line with previous findings (Billen et al., 2023), we found prominent differences in EVR expression between the stepping conditions, yet we could not identify plausible cortical correlates for proactive ‘tuning’ of the fast visuomotor network. While previous studies have implicated theta band activity in (pre)frontal cortex in proactive motor response inhibition (Adelhöfer & Beste, 2020; Wendiggensen et al., 2022), we found no evidence of such (pre)frontal involvement in EVR suppression when postural demands were high. Neither did the frontal cluster exhibit any evident changes relative to baseline during the preparatory phase, nor did we find any relevant differences in cortical dynamics between the conditions.

## Step initiation phase

To aid interpreting the EEG findings during the step initiation phase, we first summarize the observed sequence and timing of events in the two stepping conditions. Upon target reappearance, following a brief transitory epoch from the preparatory phase, the onset of the EVR (at 110-120 ms) in stance-leg GM demarcated the first target-selective event in the anterolateral stepping condition; step onset followed at ∼350 ms. In anteromedial stepping, EVRs were generally suppressed. Consistent target-selective GM activity in the stepping leg was observed around 150ms, which resulted in APA onset at ∼160 ms; average step reaction times in this condition were much slower (∼450 ms). Hence, during the first 350 ms of the step initiation phase both feet were still on the ground in either condition, but centre of mass dynamics were distinctly different.

## Differential cortical dynamics in the SMA during step initiation

Upon target reappearance, we observed continued beta power suppression in the supplementary motor area, which was marginally stronger in anteromedial step initiation (around 100-200 ms), as well as a strong theta/alpha power increase, which was also larger during anteromedial compared to anterolateral step initiation. The theta power increase appeared to be more widespread throughout multiple cortical areas (such as frontal areas and midline parietal, see Appendix) but was most pronounced in the SMA; the concurrent alpha power increase appeared to be more localized to the SMA.

Previous EEG studies investigating cortical dynamics during whole-body movements have reported similar results over the SMA. For example, Varghese and colleagues (2016) demonstrated event-related theta and alpha synchronization that appeared to be stronger and longer lasting during APA execution prior to a lateral step compared to a simple lateral weight shift, potentially reflecting enhanced sensorimotor activation in the preparation and generation of APAs.

An alternative, and not mutually-exclusive interpretation of the greater SMA theta power when stepping under greater postural demand is its involvement in stability monitoring. Previous studies have demonstrated midfrontal theta power increments when stability was challenged by external balance perturbations (Stokkermans et al., 2023) or when walking on uneven terrain (Liu et al., 2024; Sipp et al., 2013). Although the current study did not involve any external perturbations, but instead employed a global manipulation of postural demands, the theta power increase in the current study may also reflect more extensive performance monitoring during anteromedial stepping as the increased postural demands may potentially endanger balance.

In addition to its purported role in APA generation, the SMA has also been implicated in the inhibitory control of voluntary action, especially during motor tasks involving rapid choices between competing motor responses. Proper inhibition of inadequate motor responses in favour of the chosen motor response is therefore essential. Lesion studies in humans have shown that this response inhibition involves the SMA (Sumner et al., 2007). Furthermore, recordings of local field potentials in monkey SMA revealed a power increase in low-frequency bands (5-20 Hz) during and following a successful inhibition of arm movements (Stuphorn & Emeric, 2012). As our experimental paradigm involved suppression of EVRs in anteromedial steps, we speculate that the observed differential SMA dynamics may be involved in reactive inhibition of EVR responses. This inhibition may take place at the level of the midbrain reticular formation, a structure that receives projections from the SMA (Jürgens, 1984) and is thought to be involved in the modulation of EVRs (Contemori et al., 2023).

## M1 signatures of target-selective APA recruitment

We observed stronger alpha/beta I power suppression in the primary motor cortex for the anteromedial compared to the anterolateral stepping condition, with this difference between conditions being already present at the instant of target reappearance, likely as a continuation of differences observed in the late preparatory phase. As oscillatory dynamics in either condition did not differ between steps taken with the left or the right leg until ∼150 ms after stimulus reappearance (see Figure 7B), this initial contrast between postural conditions – albeit modest in strength – likely reflects non-target-selective activity for orchestrating generic motor actions in anteromedial step initiation. A strong alpha and beta I power suppression then emerged around 150-200ms, earlier and significantly stronger in anteromedial than anterolateral step initiation, indicating greater M1 involvement when initiating a step under more posturally-demanding conditions. The contrast remained significant until ∼350ms, which roughly corresponded to the time of anterolateral step onset. As during this epoch both feet were still on the ground in either condition, but with distinctly different ground reaction force profiles measured underneath both legs (see Figure 2), the execution of the APA during anteromedial step initiation likely involves an important contribution of M1.

Starting around 150ms after stimulus reappearance, the first leg-specific changes in M1 dynamics emerged according to the (prospective) stepping side. In the anterolateral stepping condition, this timing evidently lags the target-selective EVR recruitment observed in GM, supporting the purportedly subcortical origin of this earliest target-selective event. In this condition, alpha and beta suppression was more pronounced for steps with the leg contralateral to the M1 cluster, suggesting stronger cortical involvement for the stepping leg compared to the stance leg, consistent with prior studies (Gwin et al., 2011; Nordin et al., 2019).

In anteromedial stepping, the contrast between stance and stepping side was also significant, but not as pronounced and largely restricted to alpha and low beta frequencies. These findings suggest greater M1 involvement in stance leg recruitment during APA execution than during rapid anterolateral step initiation in the absence of an APA, despite ground reaction forces under the stance leg in the latter stepping condition being *higher* during the entire epoch (see Figure 2).

## Enhanced parietal engagement during APA execution

In parietal and occipital cortex, we also observed significantly stronger alpha/beta power suppression during the initiation of an anteromedial step compared to an anterolateral step. As our task involves a goal-directed stepping movement, it follows that the role of vision is crucial to guide the stepping leg towards the direction of the target. The dorsal visual stream (‘vision-for-action’ pathway) projects from visual areas towards the posterior parietal cortex and processes information for guiding goal-directed limb movements towards the desired position (Andersen & Buneo, 2003; Fattori et al., 2010; Goodale, 2011; Goodale & Westwood, 2004; Mishkin et al., 1983; Sakata, 2003). In the anterolateral condition, our findings may thus reflect less extensive visuospatial integration in parietal areas, since step initiation in this condition appears to rely on more reflexive, directly stimulus-driven visuomotor transformations at the subcortical level. Anteromedial stepping, on the other hand, requires a more voluntary and deliberate step initiation due to the increased postural demands, perhaps resulting in stronger cortical processing in parietal areas. These findings are in line with prior research (Liu et al., 2024), indicating sustained alpha and beta power suppression in the posterior parietal cortex during walking on an uneven surface compared to an even surface. Such contribution of the parietal cortex is also supported by the work of Spedden and colleagues (2022), who revealed reduced corticocortical coherence in alpha and beta/gamma between posterior parietal cortex and dorsolateral premotor cortex during visually guided step initiation and execution versus the control condition (standing and watching).

## Limitations & methodological considerations

A limitation of our study is the use of a template head model, which may lead to source-localization errors up to 2 cm compared to an anatomically accurate individual-specific head model (Liu et al., 2023). This limits interpretating the observed individual clusters in the parietal/occipital areas, which were found in close proximity. Furthermore, due to the blocked nature of the task, participants knew in advance whether the target would appear anterolaterally or anteromedially. As the postural demands are oftentimes more unpredictable in daily life, experimentally increasing the uncertainty regarding the target’s spatial location (and consequently the postural demands) may provide valuable insights in future studies. We expect that in situations where postural demands are unknown, EVR suppression and APA expression may be stronger, as part of a “default” state to prioritize postural control before executing the step itself (Castellote et al., 2024; Piscitelli et al., 2017).

## Conclusions

We investigated the cortical dynamics underlying the initiation of rapid steps during two contrasting postural demand conditions. Our collective findings point at an overall greater cortical involvement during the preparation and initiation of a visually guided step requiring an APA, as compared to a more reflexive, stimulus-driven response in the absence of an APA. Furthermore, we found little evidence of cortical activity potentially involved in context-dependent modulation of EVR expression. Instead, we speculate that APA-related cortical activity from SMA may have indirectly suppressed EVRs in the high postural demand condition, perhaps via gating mechanisms at the level of the brainstem involving the reticular formation (Contemori et al., 2023). Our findings could have functional implications for populations with impaired postural control due to aging or disease. When postural control is compromised, stepping actions that involve APAs may demand even more allocation of cortical resources compared to the young individuals studied here. If even “simple” stepping movements require more cortical engagement, it could limit the ability to flexibly adapt and interact with our dynamic environments. This is evident in reduced dual-tasking ability (Beurskens & Bock, 2012) and lower propensity to step reflexively in elderly participants and in people with PD (Billen et al., 2024). To shed more light on the impact of aging and disease on the neural control of this highly common daily-life task, future research may build upon the experimental methodology presented here, by studying modulation in cortical activity during step initiation under varying postural demands.

## Author Contributions

*Ilse Giesbers:* Conceptualization, Methodology, Software, Formal analysis, Investigation, Data curation, Writing – Original Draft, Review & Editing, Visualization; *Lucas Billen:* Conceptualization, Methodology, Software, Formal analysis, Investigation, Data curation, Writing – Original Draft, Review & Editing, Visualization; *Joris van der Cruijsen:* Methodology, Formal analysis, Writing - Review *Brian Corneil:* Conceptualization, Methodology, Software, Resources, Writing - Review & Editing, Supervision, Funding acquisition; *Vivian Weerdesteyn:* Conceptualization, Methodology, Resources, Writing - Review & Editing, Supervision, Project administration, Funding acquisition

## Funding

This work was supported by a Donders Centre for Medical Neuroscience (DCMN) grant to BDC and VW.

## Data availability statement

The authors confirm that the data supporting the findings of this study are available within the article and its supplementary materials.

## Appendix

**Figure S1.**
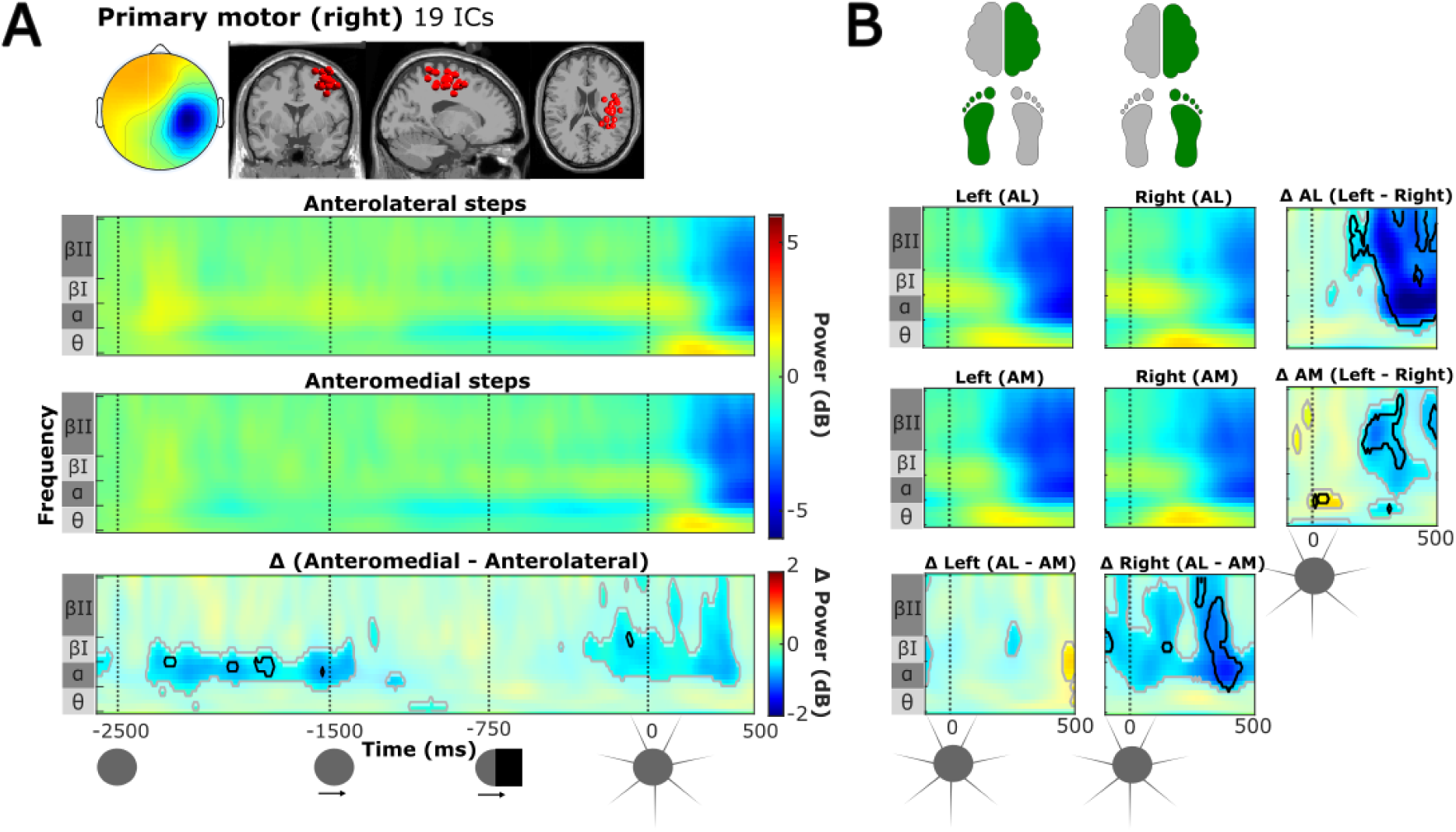
**A**. Scalp map and locations equivalent current dipoles (grey underlay), ERSPs for anterolateral (first row) and anteromedial steps (second row) and contrast plot (third row) for the right primary motor cluster. **B.** ERSPs during the step initiation phase (from target reappearance t = 0) for left and right steps during anterolateral stepping and anteromedial stepping or the right primary motor cluster, and its contrast maps.

**Figure S2.**
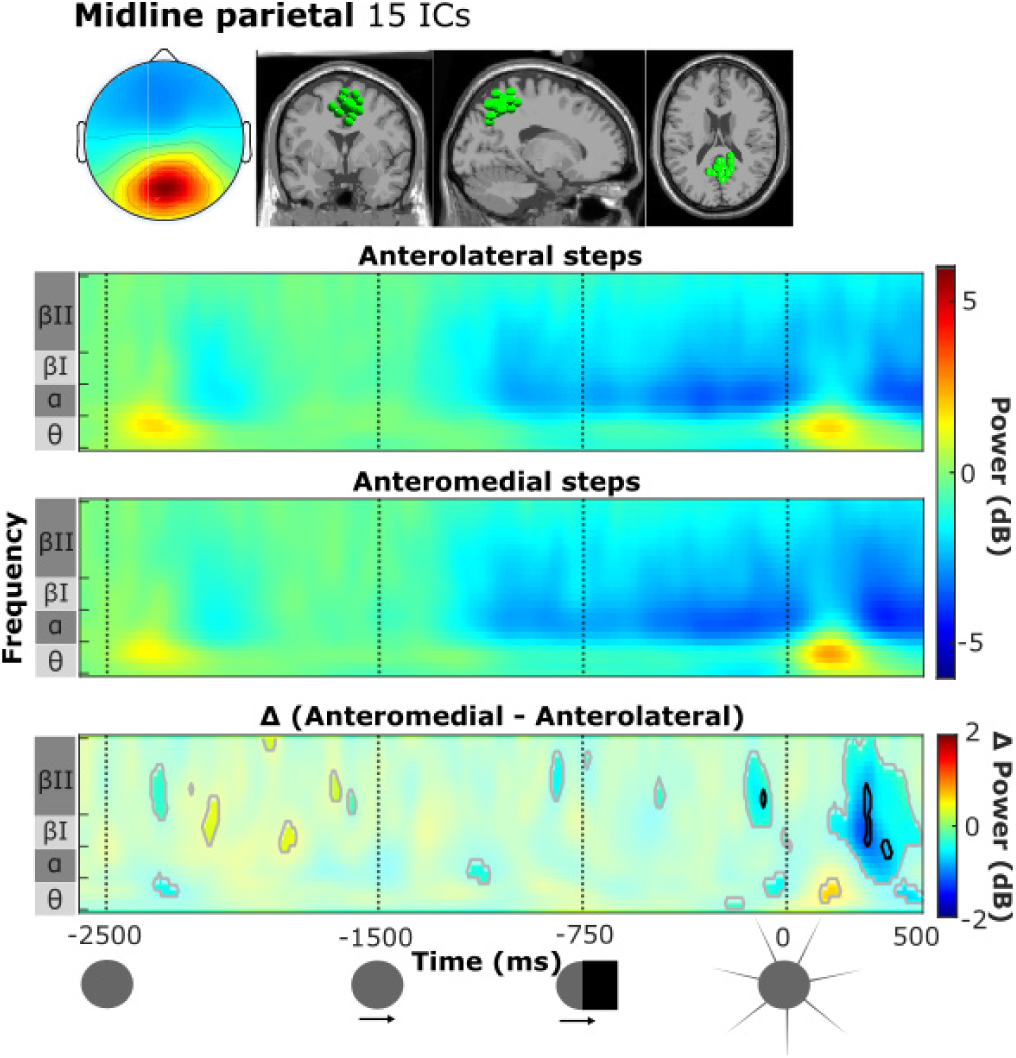
Scalp map and locations equivalent current dipoles (grey underlay), ERSPs for anterolateral (first row) and anteromedial steps (second row) and contrast plot (third row) for the midline parietal cluster.

**Figure S3.**
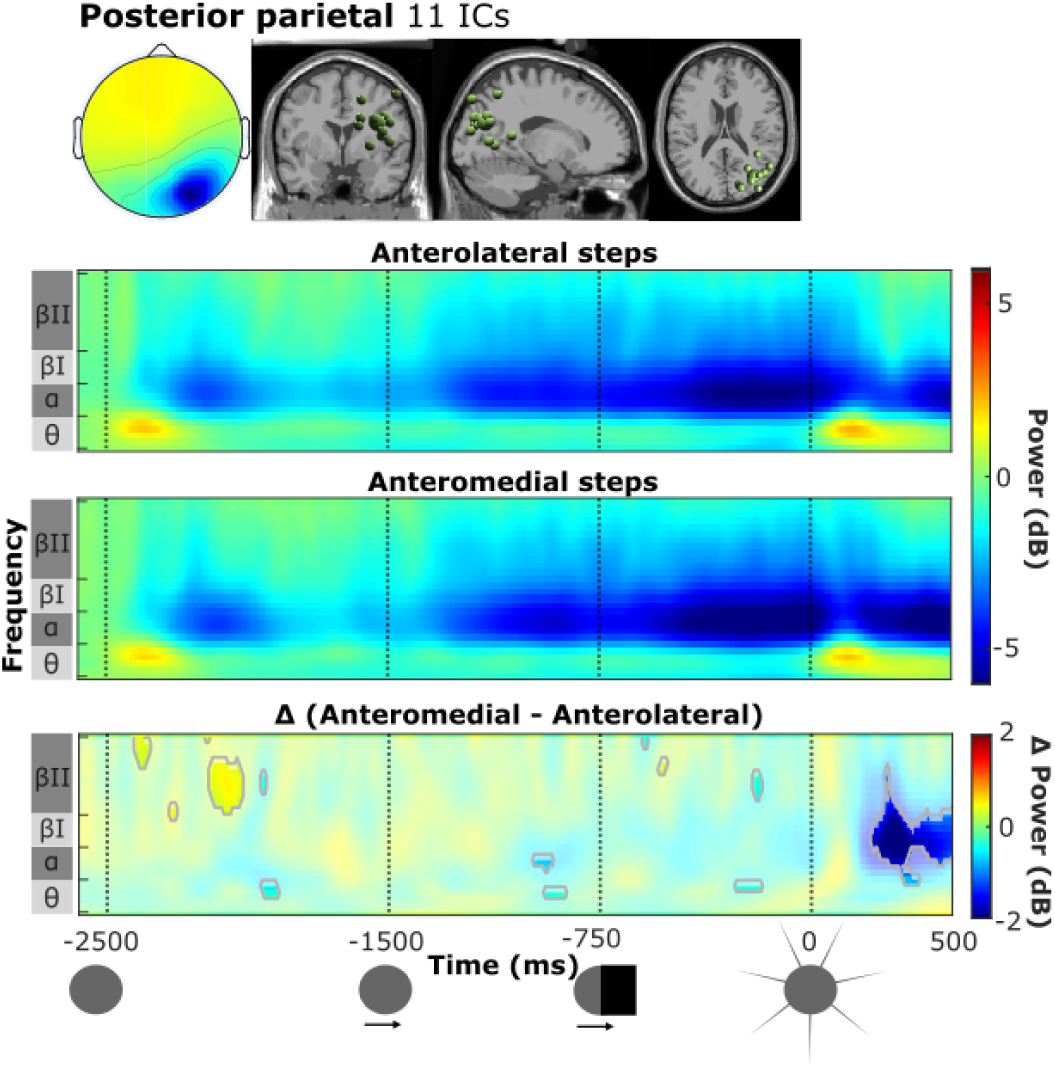
Scalp map and locations equivalent current dipoles (grey underlay), ERSPs for anterolateral (first row) and anteromedial steps (second row) and contrast plot (third row) for the posterior parietal cluster.

**Figure S4.**
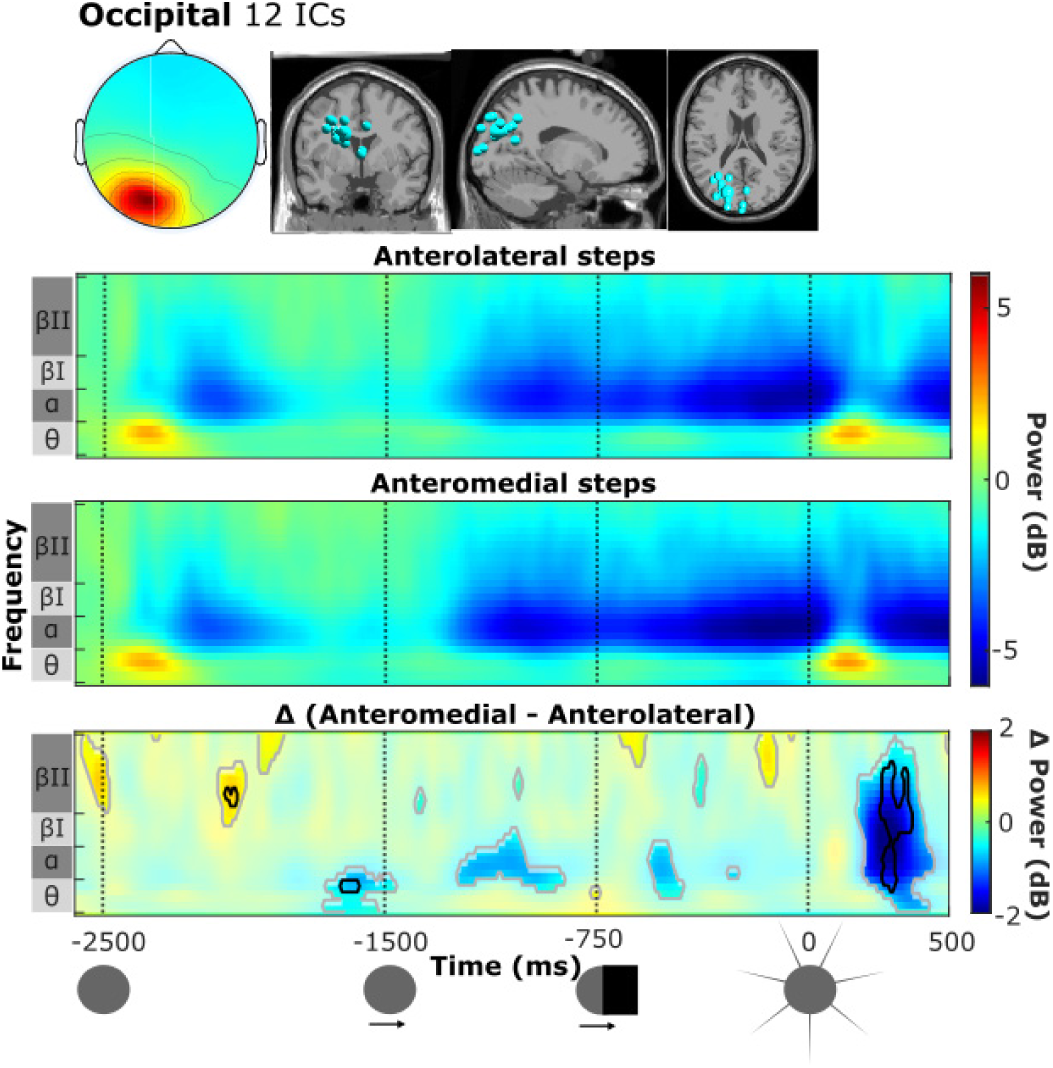
Scalp map and locations equivalent current dipoles (grey underlay), ERSPs for anterolateral (first row) and anteromedial steps (second row) and contrast plot (third row) for the occipital

## Notes

### Competing Interest Statement

The authors have declared no competing interest.

